# Serotonin signaling modulates growth and motility in juvenile *Fasciola hepatica*

**DOI:** 10.1101/2025.10.02.679961

**Authors:** Emily Robb, Sarah Muise, Lana Watt, Rebecca Armstrong, Duncan Wells, Paul McCusker, John Harrington, Andreas Krasky, Paul M. Selzer, Nikki J. Marks, Aaron G. Maule

**Affiliations:** Understanding Health & Disease, School of Biological Sciences, Queen’s University Belfast, Belfast BT9 5DL, UK; Boehringer Ingelheim Animal Health, 1730 Olympic Dr, Athens, GA 30601, USA; Boehringer Ingelheim Animal Health, Binger Str. 173, 55216 Ingelheim am Rhein, Germany

## Abstract

*Fasciola hepatica* causes fasciolosis, a parasitic disease that poses significant animal and human health challenges. Control relies on flukicides, most of which are adulticides, with only triclabendazole effective against the pathogenic migratory juvenile. Classical neurotransmitter pathways are widely targeted by anthelmintics yet remain underexplored for flukicide development. Here we explore the importance of serotonin (5-HT) signaling in juvenile fluke. *In silico* analyses confirmed all *F. hepatica* life stages express a complete 5-HT signaling pathway encompassing genes encoding proteins for 5-HT synthesis, transport, and reuptake, as well as five putative 5-HT G protein-coupled receptors (GPCRs). Homology and binding motif analyses supported the presence of two 5-HT_1_ (Fh5HT_1A_, Fh5HT_1B_) and three 5-HT_7_ (Fh5HT_7A_, _-7B_, _-7C_) GPCRs. Immunocytochemistry and *in situ* hybridization revealed widespread neuronal expression of 5-HT, its synthetic enzyme tryptophan hydroxylase (FhTPH), and the GPCR Fh5HT_7C_. 5-HT addition stimulated juvenile fluke motility; consistent with this observation, serotonin reuptake inhibition, which causes 5-HT persistence at synaptic junctions, also enhanced juvenile movement. Silencing of FhTPH, a key enzyme in 5-HT synthesis, blunted juvenile motility, a phenotype reversed by the addition of 5-HT. Silencing the fluke vesicular monoamine transporter (FhVMAT), which packages 5-HT into synaptic vesicles, reduced juvenile motility, whilst silencing the 5-HT reuptake transporter (FhSERT) which recycles synaptic 5-HT increased juvenile motility and growth, consistent with 5-HT accumulation enhancing effects. Whilst combinatorial silencing of Fh5HT_1_ receptors reduced fluke motility, silencing Fh5HT_7_ receptors led to a greater reduction in motility. Exogenous addition of 5-HT partially rescued motility deficits of juveniles with silenced Fh5HT_1_ receptors, but 5-HT excitation was abolished in Fh5HT_7_-RNAi juveniles, exposing their importance to fluke motility. Notably, sustained 5-HT exposure promoted juvenile growth, but these effects were not blunted by receptor-RNAi. The findings emphasize a central role of serotonin signaling in both juvenile motility and growth, exposing novel aspects of receptor function and encouraging therapeutic exploitation for liver fluke control.

**Author Summary:** The liver fluke, *Fasciola hepatica*, causes fasciolosis, a neglected tropical disease that poses a significant burden on human and animal health. There is no vaccine for fasciolosis and treatment relies on a single drug, triclabendazole, to control the early stages of infection which cause liver pathology whilst migrating through the mammalian host. Single drug reliance has increased the incidence of drug resistance in both human and animal populations, such that there is a pressing need for the characterization of novel drug targets and development of new anthelmintics targeting liver fluke. The focus of this research is to examine the role of the serotonin signaling system of liver fluke, bridging a gap in knowledge to enable the exploitation of this signaling pathway for flatworm drug development. Here, bioinformatic analysis has characterized the pathway components and receptors in multiple clinically relevant flatworm parasite species. Chemical and functional genomic methods have been used to prove the integral function of serotonin in liver fluke biology, regulating motility and growth, both essential for parasite infection and survival. This work provides data that help validate the serotonergic system of liver fluke as a potential target for future anthelmintic development.

## Introduction

Fasciolosis, caused by the liver fluke *Fasciola hepatica*, is a global issue affecting both livestock and humans, with agricultural losses estimated at $3.2 billion annually (1) . As a neglected tropical disease, it impacts approximately 17 million people in some of the world’s poorest regions. The most effective treatment for fasciolosis is the anthelmintic drug triclabendazole (2). Triclabendazole is the only drug effective against the early juvenile stages of the parasite, which cause the most severe damage during their migration through the hosts liver (3). However, the heavy reliance on triclabendazole for treatment has led to increasing cases of drug resistance in both human and livestock populations, underscoring the need for the development of new drugs to combat these parasites (4,5).

The flatworm nervous system has well developed classical neurotransmitter and neuropeptidergic systems, which are integral to regulating biological activities essential for parasite survival, including motility, reproduction, growth, and development (6). Research has identified these systems as a source of potential druggable targets. In particular, the neuropeptidergic system has attracted considerable attention, with numerous studies reporting drug target validation from within this pathway (6,7). Notably, neurotransmitter pathways have been successfully targeted in nematode parasites, with highly effective anthelmintic drugs such as levamisole and ivermectin (8). However, the relative sparsity of studies characterizing analogous flatworm neurotransmitter pathways has hindered similar progress in drug development for flatworm parasites (9)

Serotonin (5-HT), an ancient signaling molecule conserved across metazoans, is one of the most extensively studied neurotransmitters, particularly in higher animals and humans. Its roles extend beyond modulating neuronal activity to include the regulation of processes such as metabolism and immune responses (10,11). In vertebrates, endogenous 5-HT synthesis occurs through a well-characterized pathway involving the enzymes tryptophan hydroxylase (TPH) and aromatic L-amino acid decarboxylase (AADC), which convert tryptophan into 5-HT (12,13). Once synthesized, 5-HT is packaged into synaptic vesicles by vesicular monoamine transporters (VMAT) and released into the synaptic cleft (12,13). 5-HT signals via a 5-HT-gated cation-selective ion channel (5-HT3) and/or G protein coupled receptors (GPCRs; 5HTR) resulting in an increase or decrease in cyclic adenosine monophosphate (cAMP) second messenger production by adenylate cyclase (12,13). 5-HT signaling is terminated by 5-HT reuptake transporters (SERT), which clear the neurotransmitter from the synapse (12,13). In parasitic helminths, bioinformatic studies suggest that a 5-HT synthesis and signaling pathway similar to that of vertebrates is conserved. However, while GPCR-mediated 5-HT signaling has been characterized in helminths, a 5-HT-gated ion channel specific to flatworm parasites has yet to be identified (14,15). Parasitic flatworms have also been shown to actively transport exogenous 5-HT from hosts across their tegument further suggesting functional importance for parasite survival within a host (16–18).

Despite an extensive history of study, the function of 5-HT in flatworm biology remains poorly characterized for drug target exploitation. Early studies show high expression of 5-HT in both the central and peripheral nervous system with prevalence in neurons linked with innervation of the digestive tract and body wall muscles of adult *F. hepatica* (19,20). Rich serotonin immunoreactivity was also associated with the oral and ventral suckers and surrounding nerve fibres of *F. hepatica* (19). Evidence strongly supports serotonin’s role in modulating neuromuscular function and motility in flatworms. For instance, 5-HT excites muscle strips in liver fluke (21,22), as well as in other parasitic flatworms such as the cestode *Hymenolepis diminuta* (23) and monogenean *Diclidophora merlangi* (24). Whole-worm motility assays further underscore 5-HT’s role in modulating flatworm motility, as it stimulates larval motility in the blood fluke *Schistosoma mansoni* (25) and other flatworm species (26,27). Direct actions on muscle were supported by the fact that 5-HT was found to be critical for dispersed muscle fiber contraction in *S. mansoni* (28,29). Molecular studies have further corroborated serotonin’s importance as a regulator of parasite movement. For example, RNA interference (RNAi)-based silencing of the 5-HT reuptake transporter (smSERT) in *S. mansoni* led to a marked increase in schistosomula motility, while knockdown of a serotonin receptor (sm5HTR) resulted in a significant decrease in motility (18,30). The role of 5-HT in modulating flatworm motility encourages consideration as a therapeutic target in parasitic flatworms.

Beyond its role in motility, 5-HT has been identified as a metabolic regulator in parasitic flatworms, enhancing carbohydrate metabolism and facilitating glucose uptake to increase ATP availability, thereby supporting energy-intensive processes such as muscle contraction (31–33). Emerging evidence suggests 5-HT also plays a significant role in regulating cell proliferation and development, indicating a crucial non-neuronal function. In mammals, deficiencies in 5-HT signaling during embryogenesis result in morphogenetic abnormalities, emphasizing its importance as a developmental regulator (34). Additionally, 5-HT receptor antagonists have demonstrated anticancer properties in several *in vitro* studies, further supporting serotonin’s involvement in cell growth and proliferation (35). In planarians, 5-HT is essential for eye regeneration in *Schmidtea mediterranea*, showcasing its role in tissue repair and developmental processes (36). In parasitic flatworms, 5-HT has similarly been implicated in development, for example it stimulated the development of *Echinococcus multilocularis* metacestodes and induced cell proliferation, suggesting its importance in cestode growth and survival (37). This multifunctionality in regulating metabolic, motility and developmental pathways encourages evaluation of 5-HT signaling as a therapeutic target for parasite control.

There is substantial evidence to support the hypothesis that 5-HT signaling is critical to a variety of essential systems in flatworms. Coupled with pharmacological studies indicating the druggability of parasite-specific 5-HT receptors (38), this signaling system presents a promising avenue for the identification of novel therapeutic targets. However, critical knowledge gaps persist, as the role and importance of multiple components of the 5-HT signaling systems in flatworms remain poorly characterized, impeding drug development efforts.

This study aims to characterize the 5-HT signaling pathway for the liver fluke, *F. hepatica*. Functional genomics complemented by expression analyses and the chemical validation of 5-HT pathway components underscore their critical role in regulating the motility of the juvenile liver fluke, whilst simultaneously exposing a broader role for serotonin in parasite growth. Additionally, bioinformatic analyses identified 5-HT-gated G protein-coupled receptors (GPCRs) across multiple clinically relevant flatworm species, encouraging exploitation of this pathway for the development of novel flukicides/anthelmintics.

## Results and Discussion

### *F. hepatica* juveniles express a complete gene set for 5-HT signaling

*In silico* analysis (Figure 1A) revealed significant conservation of 10 genes linked to 5-HT signaling in *F. hepatica.* These include five genes encoding proteins involved in 5-HT synthesis, transport, and reuptake: a tryptophan hydroxylase (FhTPH); two aromatic L-amino acid decarboxylases (FhAADC); a vesicular monoamine transporter (FhVMAT); and a 5-HT reuptake transporter (FhSERT) (S1 Table). Additionally, five putative serotonin G protein-coupled receptor (GPCR) genes (Fh5HTR) were identified (Figure 1 B&C; S1 Table). All identified genes exhibit high conservation of functional domains, as observed in orthologues from humans (39–42) and blood fluke, *Schistosoma mansoni* (Figure 1D) (15,18,30,43). Notably, *F. hepatica* expresses two distinct AADC enzymes (FhAADC1 & FhAADC2), whereas humans possess only one. This gene duplication, which has also been observed in other flatworm species, may reflect an additional role in dopamine signaling (15). These findings align with genomic studies demonstrating conservation of 5-HT signaling components across diverse flatworm species (15,44). Transcriptome analysis revealed peak expression of 5-HT signaling pathway genes during the newly excysted juvenile (NEJ) stage, suggesting a role for 5-HT in the highly motile infective and migratory phases of the parasite’s life cycle (Figure 1E and S1 Table), consistent with the hypothesis that 5-HT is essential for neuromuscular function. Interestingly, the synthesis enzymes FhTPH and FhAADC also show elevated expression during the egg stage, supporting a non-neuronal role for 5-HT in egg development and biology (Figure 1E and S1 Table). This observation suggests an important role for 5-HT in the egg stage, where the assimilation of exogenous 5-HT is likely limited, necessitating endogenous production to fulfil developmental requirements. Five putative 5-HT-GPCRs were identified in *F. hepatica;* FhHiC23_g2340, FhHiC23_g7880, FhHiC23_g4917, FhHiC23_g1196 and FhHiC23_g115 (Figure 1B and S1 Table). Consistent with the literature, a 5-HT-gated ion channel was not identified as part of this analysis (15). Alignments with characterized human 5HT_1_ (Q5ZGX3) and functionally characterized *S. mansoni* 5HT_7_ (Smp_126730; (30)) 5-HT-gated GPCRs show conservation of key motifs essential for ligand binding (DVXXCT and WXXF, where X denotes any amino acid) and receptor activation (DRY) (45) (Figure 2A).

**Figure 1.**
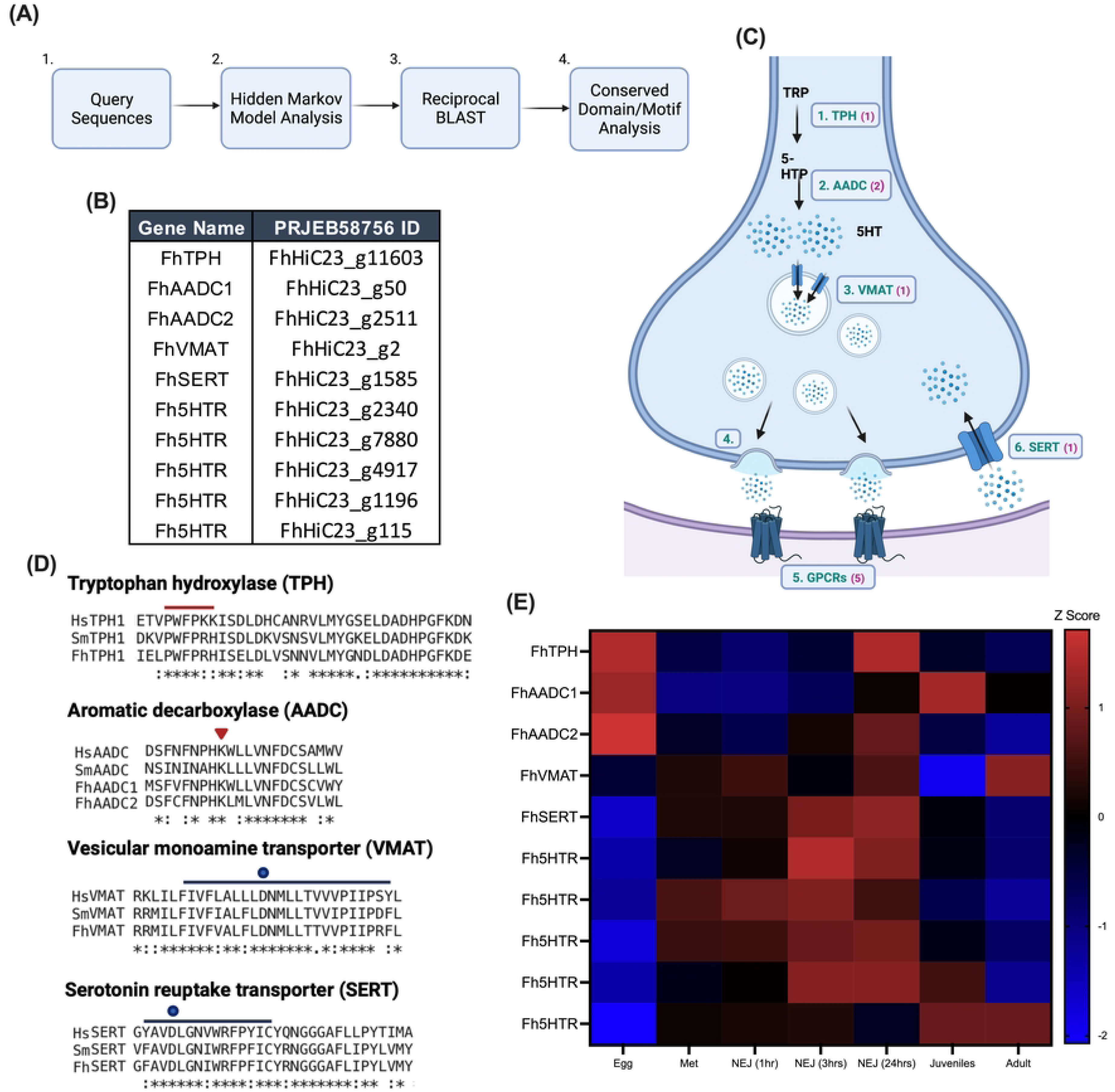
*Fasciola hepatica* possesses serotonin signaling pathway genes. **(A) Schematic representation of the bioinformatic pipeline used to identify serotonin signaling genes in *F. hepatica*. (B) Gene identifiers (IDs) of serotonin-related components identified in the *F. hepatica* genome assembly PRJEB58756.** Abbreviations: FhTPH = tryptophan hydroxylase; FhAADC = aromatic L-amino acid decarboxylase; FhVMAT = vesicular monoamine transporter; FhSERT = serotonin reuptake transporter; Fh5HTR = serotonin-gated G protein-coupled receptors (GPCRs). **(C) Illustration of serotonin signaling at a synapse.** (1) The precursor amino acid tryptophan (TRP) is converted to 5-hydroxytryptophan (5-HTP) by tryptophan hydroxylase (TPH). (2) 5-HTP is further converted to serotonin (5-hydroxytryptamine, 5-HT) by aromatic L-amino acid decarboxylase (AADC). (3) 5-HT is sequestered into synaptic vesicles via vesicular monoamine transporters (VMAT). (4) Vesicles fuse with the synaptic membrane, releasing 5-HT into the synaptic cleft. (5) Released 5-HT binds to GPCRs on the postsynaptic neuron to propagate signaling. Note: although serotonin can also interact with ligand-gated ion channels (LGICs), no 5-HT-sensitive LGICs were identified in the *F. hepatica* genome. Magenta-bracketed numbers indicate the number of encoding genes identified in genome PRJEB58756. **(D) Sequence alignments of *F. hepatica* serotonin signaling genes with homologous sequences from *Homo sapiens* (Hs-) and *Schistosoma mansoni* (Sm-).** Red lines highlight conserved catalytic domains in TPH, and red arrowhead indicates pyridoxal-phosphate binding site in AADC. Blue lines represent transmembrane domain 1 of VMAT and SERT, while a blue dot marks a key aspartic acid residue implicated in ligand binding. (**E) Heatmap displaying expression levels of *F. hepatica* serotonin signaling genes across different life cycle stages (egg to adult).** Z-scores were calculated from transcripts per million (TPM) data obtained from WormBase ParaSite (release 54) and generated by Cwiklinski et al. (46).

**Figure 2.**
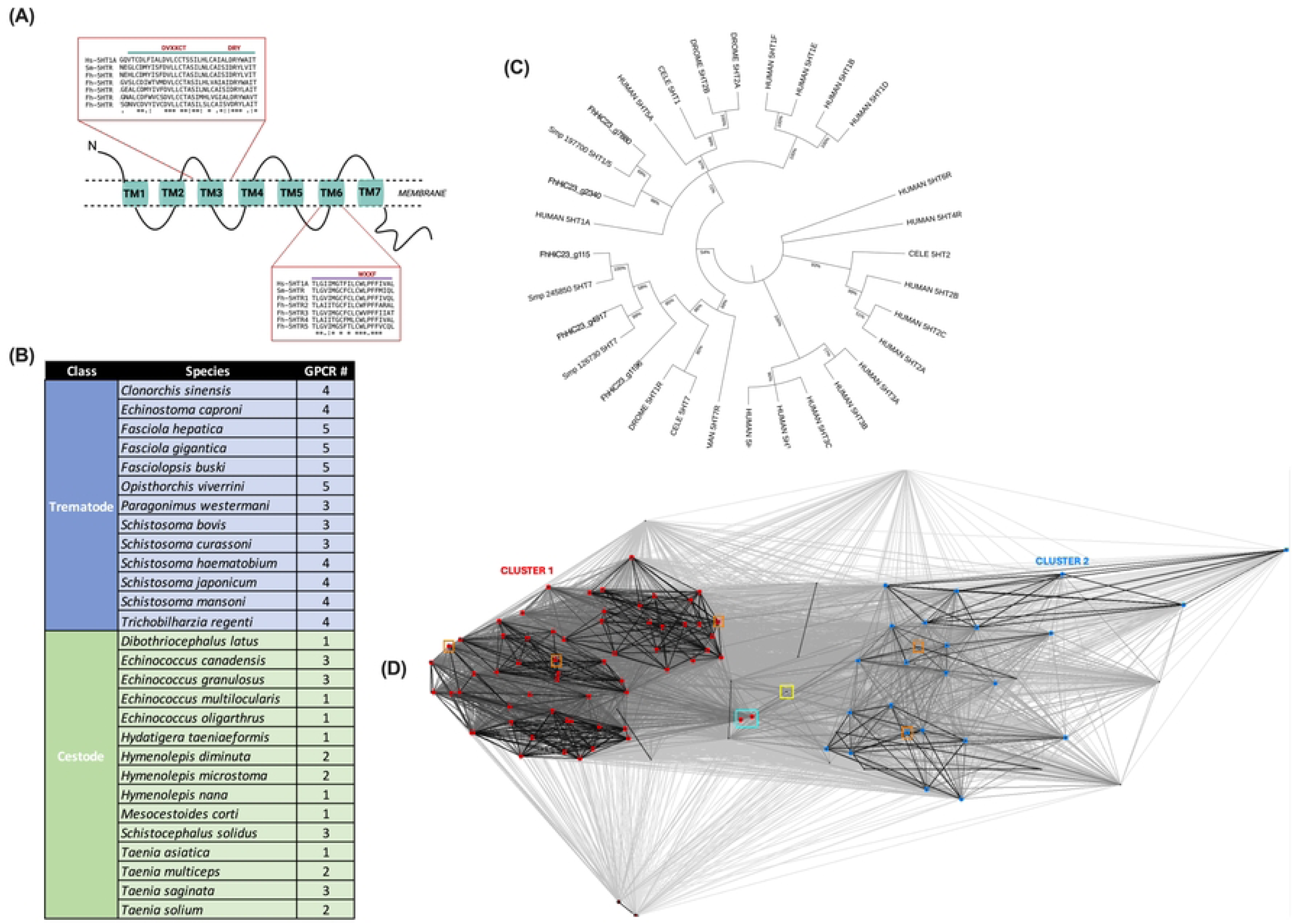
Comparative and phylogenetic analysis of serotonin-gated G protein-coupled receptors (GPCRs) in *Fasciola hepatica* and other parasitic flatworms. **(A) Multiple sequence alignment of *F. hepatica* serotonin GPCRs with homologous sequences from *Homo sapiens* (Hs-) and *Schistosoma mansoni* (Sm-).** The green line highlights transmembrane domain three (TM3), while the purple line highlights transmembrane domain six (TM6). Conserved serotonin-binding motifs (DVXXCT and WXXF) essential for ligand recognition, as well as the canonical ‘DRY’ motif critical for GPCR activation, are outlined in red. **(B) Summary table displaying the number of serotonin-gated GPCRs identified across clinically relevant trematode and cestode species**. Genomic data were obtained from WormBase ParaSite (version 19) and analyzed for the presence of serotonin GPCR homologs. The species investigated include *Echinococcus canadensis* (PRJEB8992), *Echinococcus granulosus* (PRJNA182977 - G1), *Echinococcus multilocularis* (PRJEB122 - Java), *Echinococcus oligarthrus* (PRJEB31222 - DaMi1*), Hymenolepis diminuta* (PRJEB507 - Denmark), *Hymenolepis microstoma* (PRJEB124), *Hymenolepis nana* (PRJEB508 - Japan), *Mesocestoides corti* (PRJEB510 - Specht & Voge 1965), *Taenia asiatica* (PRJEB532 - South Korea), *Taenia multiceps* (PRJNA307624 - Gns01), *Taenia saginata* (PRJNA71493 - TSAYD01), *Taenia solium* (PRJNA170813 - Mexico), *Dibothriocephalus latus* (PRJEB1206*), Spirometra erinaceieuropaei* (PRJEB1202), *Schistocephalus solidus* (PRJEB527 - NST_G2), *Hydatigera taeniaeformis* (PRJEB534 - Spain/Canary Islands), *Clonorchis sinensis* (PRJNA386618 - Cs-k2), *Schistosoma bovis* (PRJNA451066), *Schistosoma japonicum* (PRJNA520774 - HuSjv2), *Fasciolopsis buski* (PRJNA284521 - HT), *Fasciola gigantica* (PRJNA230515 - Uganda_cow_1), *Schistosoma haematobium* (PRJNA78265), *Schistosoma curassoni* (PRJEB519 - Senegal/Dakar), *Schistosoma mansoni* (PRJEA36577*), Trichobilharzia regenti* (PRJEB44434 - tdTriRege1.1), *Opisthorchis viverrini* (PRJNA222628), *Paragonimus westermani* (PRJNA454344), and *Echinostoma caproni* (PRJEB1207 - Egypt). **(C) Maximum likelihood phylogenetic tree depicting the evolutionary relationships between *F. hepatica* serotonin-gated GPCRs and homologous GPCR sequences from other species.** Reference sequences were obtained from *Homo sapiens* (5-HT1A-F, 5-HT2A-C, 5-HT3A-E, 5-HT5A, and 5-HT7), *Caenorhabditis elegans* (5-HT1, 5-HT2, and 5-HT7), and *Drosophila melanogaster* (5-HT1 and 5-HT2A/B) through UniProt (53). Additionally, published *S. mansoni* GPCR sequences (Smp_197700, Smp_245850, and Smp_126730) were included based on prior literature (references 14, 29, 55). **(D) CLANS (Cluster Analysis of Sequences) diagram visualizing the clustering relationships of serotonin-gated GPCRs from all examined parasitic flatworm species.** Clustering groups are color-coded based on sequence similarity and evolutionary relationships. Red shows flatworm 5-HT GPCR cluster 1 sequences; blue shows flatworm 5-HT GPCR cluster 2 sequences; orange boxes show clustering of *F. hepatica* 5-HT GPCR sequences; yellow box shows human 5-HT_1_ GPCR sequence; cyan box shows *D. melanogaster* 5-HT_2_ GPCR sequences.

### 5-HT-gated GPCRs are conserved across clinically relevant flatworms

A further 75 flatworm 5-HT-gated GPCRs were identified across clinically significant cestode and trematode species (S2 Table), representing the most extensive dataset of flatworm 5-HT receptors to date. All 75 identified flatworm GPCRs exhibit conservation of 5-HT-associated ligand binding and receptor activation motifs, as previously characterized for *F. hepatica* (S1 Figure). Notably, a greater number of 5-HT receptors were found in trematode species (average = 4) compared to cestode species (average = 2) (Figure 2B). Interestingly, Wheeler et al. (47) reported that the size of chemoreceptor families was found to correlate with the presence of environmental and extra-host stages in the life cycles of nematodes. It is possible that the number of 5-HT receptors correlates with parasite lifestyle, however, this requires further investigation. Recently, Camicia et al. (48) presented evidence suggesting that cestode 5HT_1_ receptors may have evolved distinct motif patterns, potentially making them harder to detect using bioinformatic approaches. As a result, it is possible that some sequences may have been excluded from this analysis.

### *F. hepatica* express 5HT_1_ and 5HT_7_ type GPCRs

Current understanding of 5-HT receptors in parasitic flatworms has been mostly driven by *in silico* approaches given the limited availability of functional data for receptor characterization (49). As a result, receptor classification has largely been conducted through homology-based approaches, comparing flatworm receptors to their vertebrate counterparts (50). Through such methods, most 5-HT receptors identified in parasitic flatworms have been classified into the 5-HT_1_ and 5-HT_7_ clades (15,30,48,51). Functionally, 5HT_1_ receptors couple to G_i/o_ proteins to inhibit adenylate cyclase and decrease cAMP, whilst activation of 5HT_7_ receptors coupled to G_s_ proteins activate adenylate cyclase and increase intracellular cAMP (48).

Phylogenetic analysis grouped two *F. hepatica* 5-HT-gated GPCRs, FhHiC23_g2340 and FhHiC23_g7880, alongside human and *S. mansoni* 5HT_1_ receptors, previously identified via *in silico* analysis (Figure 2C) (15). Three further *F. hepatica* 5-HT-activated GPCRs, FhHiC23_g4917, FhHiC23_g1196 and FhHiC23_g115, were phylogenetically associated with the human 5-HT_7_ receptor (P34969) and functionally characterized 5-HT_7_ receptor (Smp_126730) from *S. mansoni* (30) suggesting these are 5-HT_7_ family serotonin-gated GPCRs (Figure 2C). Further corroboration using CLANs and clustering analysis revealed two distinct groups of flatworm 5-HT-gated GPCRs (Figure 2D and S2 Table). Cluster 1 contains the three *F. hepatica* GPCRs tentatively assigned to the 5-HT_7_ receptor family (FhHiC23_g4917, FhHiC23_g1196, and FhHiC23_g115). This cluster shows close sequence similarity with *S. mansoni’s* functionally characterized 5-HT_7_ receptor (Smp_126730), but notably, the human 5-HT_7_ receptor did not cluster with any flatworm GPCRs (Figure 2D). This sequence divergence between human and flatworm receptors could highlight an opportunity to selectively target flatworm-specific 5-HT receptors with anti-parasitic drugs. Cluster 2 includes *F. hepatica’s* FhHiC23_g2340, identified as a likely 5HT_1_-type receptor, and shows strong clustering with *S. mansoni* 5-HT_1_ receptor genes (Smp149770 and Smp197700). Interestingly, while FhHiC23_g7880 groups more closely with 5HT_1_ receptors, it did not cluster distinctly with any characterized 5-HT receptor family in the CLANs analysis (Figure 2D). Motif analysis confirmed that all *F. hepatica* 5-HT receptors share a conserved pattern of cysteine(C) and threonine (T) residues in transmembrane 3, indicative of either the 5-HT_1_ or 5-HT_7_ receptor clades (S2 Figure) (48). Notably, FhHiC23_g2340 and FhHiC23_g7880 exhibit a conserved tryptophan (W) at position 28 within transmembrane domain 3, a defining feature of 5HT_1_ receptors, while the remaining receptors possess tyrosine (Y) at this position, aligning them with 5HT_7_ receptors (S2 Figure). Furthermore, in transmembrane domain 5, at positions 42 and 46, FhHiC23_g2340 and FhHiC23_g7880 display serine (S) and alanine (A) residues, consistent with 5-HT_1_ receptor classification (S2 Figure). In contrast, the other three receptors have alanine (A) residues at both positions, matching the 5-HT_7_ receptor profile as described by Camicia et al. (48) (S2 Figure).

The binding of 5-HT to GPCR receptors induces a conformational change, triggering the release of G proteins to propagate a signaling cascade (52). To predict the most likely G protein coupling of *F. hepatica* serotonin receptors*, in silico* analysis was performed to assess structural motifs (53) (S3 Figure). FhHiC23_g2340 and FhHiC23_g7880 possessed motifs known to have coupling specificity for Gi/o family G proteins, a hallmark of 5HT_1_ receptors, further supporting their classification as 5-HT_1_ type (S3 Figure). The three 5-HT-gated GPCRs phylogenetically grouped as 5-HT_7_ type exhibited motifs with varying levels of G protein coupling specificity (S3 Figure). FhHiC23_g4917 displayed positive coupling exclusively to Gs proteins, which are associated with 5-HT_7_ receptor signaling, supporting classification as a 5-HT_7_ receptor (S3 Figure). Although FhHiC23_g115 also showed positive coupling to Gi/o proteins, it exhibited highest specificity for Gs proteins, further validating its designation as a 5-HT_7_ receptor (S3 Figure). Interestingly, FhHiC23_g1196 showed positive coupling to all G proteins analyzed, exposing the potential for promiscuity in its interactions and biological function (S3 Figure).

Based on the analysis outlined, *F. hepatica* 5-HT GPCRs were designated as follows; FhHiC23_g2340-5HT_1AFhep_, FhHiC23_g7880-5HT_1BFhep_, FhHiC23_g4917-5HT_7AFhep_, FhHiC23_g115-5HT_7BFhep_ and FhHiC23_g1196-5HT_7CFhep_.

### Modelling of *Fasciola hepatica* 5-HT-gated G protein coupled receptors

To probe the presence of 5-HT binding sites and determine structural similarity to human homologues, *F. hepatica* 5-HT GPCRs were modelled using PyMOL (81) (Figure 3A-E). Ligand docking analysis confirmed all receptors could bind 5-HT ligand in a similar region to human 5-HT receptors (82) (Figure 3A-E and S4 Figure). Comparing binding sites of 5HT_7_ receptors highlighted consensus residues hypothesized to be key for 5-HT ligand binding (Figure 3F), including Aspartate (D), Valine (V) and Threonine (T) in transmembrane 3, Glutamine (Q), Alanine (A) and Threonine (T) in transmembrane 5 and Glutamine (G)/Alanine (A), Phenylalanine (F) in transmembrane 6 (Figure 3F). Similar analysis of 5HT_1_ receptors also highlighted the presence of key residues within transmembranes 3, 5 and 6 (Figure 3F), including Aspartate (D), Valine (V) and Cysteine (C) in transmembrane 3, Phenylalanine (F)/IsoLeucine (I) and Serine (S) in transmembrane 5 and Proline (P), Phenylalanine (F), Alanine (A) and Leucine (L) in transmembrane 6 (Figure 3F). These similarities support the importance of these residues for ligand binding in liver fluke 5-HT receptors. Alignment analysis compared the structure of *F. hepatica* 5-HT GPCRs to human 5-HT GPCRs, generating a root mean square deviation (RMSD) value of similarity - the lower the RMSD value, the higher the structural similarity between two proteins with an RMSD value of <2 Å considered highly similar. *F. hepatica* 5HT_1_ receptors had RMSD values of 0 (5HT_1BFhep_) and 0.684 (5HT_1AFhep_) which indicates high similarity - indeed an RMSD value of 0 suggests the absence of conformational changes (S4 Figure). *F. hepatica* 5HT_7_ receptors showed RMSD values of 0.684 (5HT_7AFhep_), 0.760 (5HT_7BFhep_) and 0.791 (5HT_7CFhep_), again showing structural similarity with human 5-HT GPCRs (S4 Figure).

**Figure 3.**
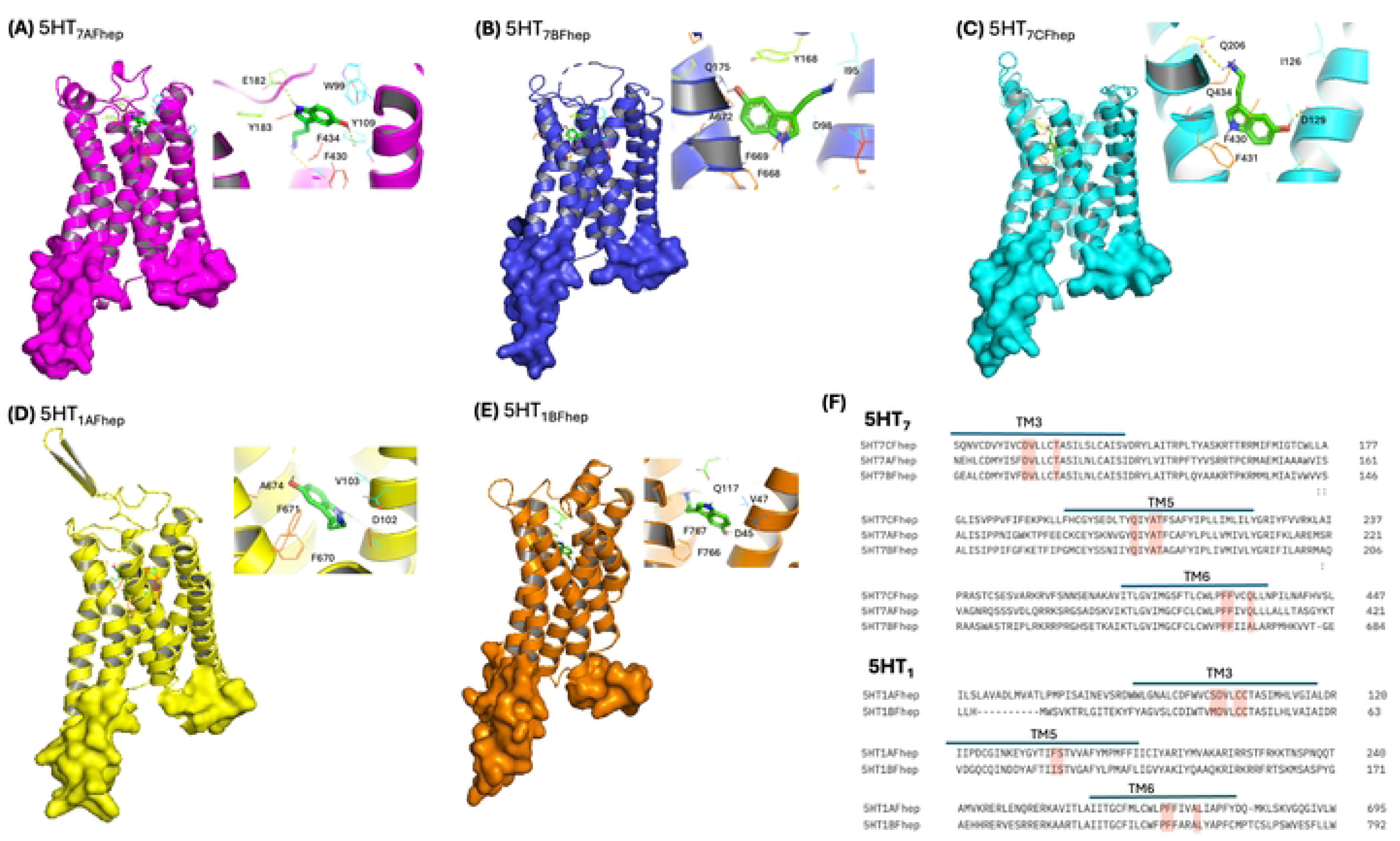
Structural models of putative *Fasciola hepatica* 5-HT-gated G protein coupled receptors (GPCRs) support the presence of residues critical to serotonin binding. **(A) Predicted structural confirmation of 5HT_7AFhep_ GPCR showing key ligand binding residues. (B) Predicted structural confirmation of 5HT_7BFhep_ GPCR showing key ligand binding residues. (C) Predicted structural confirmation of 5HT_7CFhep_ GPCR showing key ligand binding residues. (D) Predicted structural confirmation of 5HT_1AFhep_ GPCR showing key ligand binding residues. (E) Predicted structural confirmation of 5HT_1BFhep_ GPCR showing key ligand binding residues.** All structures predicted and visualized using Phyre2 (80) and PyMOL (81) software. The serotonin (SRO) ligand was imported directly from the Protein Data Bank (https://www.rcsb.org) for binding analysis carried out by DockingPie (82). **(F) Sequence alignments of *F. hepatica* 5HT_7_ and 5HT_1_ receptors showing predicted essential amino acids for ligand binding.** Ligand binding predicted by DockingPie (82). Blue line highlights transmembrane regions and red shading highlights conserved amino acids predicted by docking analysis.

### 5-HT signaling components are highly expressed in nervous system of *F. hepatica*

Immunocytochemistry demonstrates widespread 5-HT expression in the central and peripheral nervous systems of 3-week-old juvenile *F. hepatica* (5-HT expression has been reported previously in the adult and NEJ life stages), supporting its extensive role in neuromuscular and physiological regulation (Figure 4A and 4B). 5-HT immunoreactivity (IR) is prominently localized along the ventral nerve cords and in large neuronal cell bodies in and around the cerebral ganglia and associated commissures at the anterior end of the parasite, consistent with the structural organization observed in other trematodes (54) (Figure 4A-D). The IR pattern extends to the peripheral nervous system, with innervation in regions such as the pharynx and fine nerves adjacent to the surface (Figure 4A, 4B and 4D). These findings align with studies where serotonin has been implicated in controlling feeding and motility behaviors of related flatworms (30,37,51,55,56).

**Figure 4.**
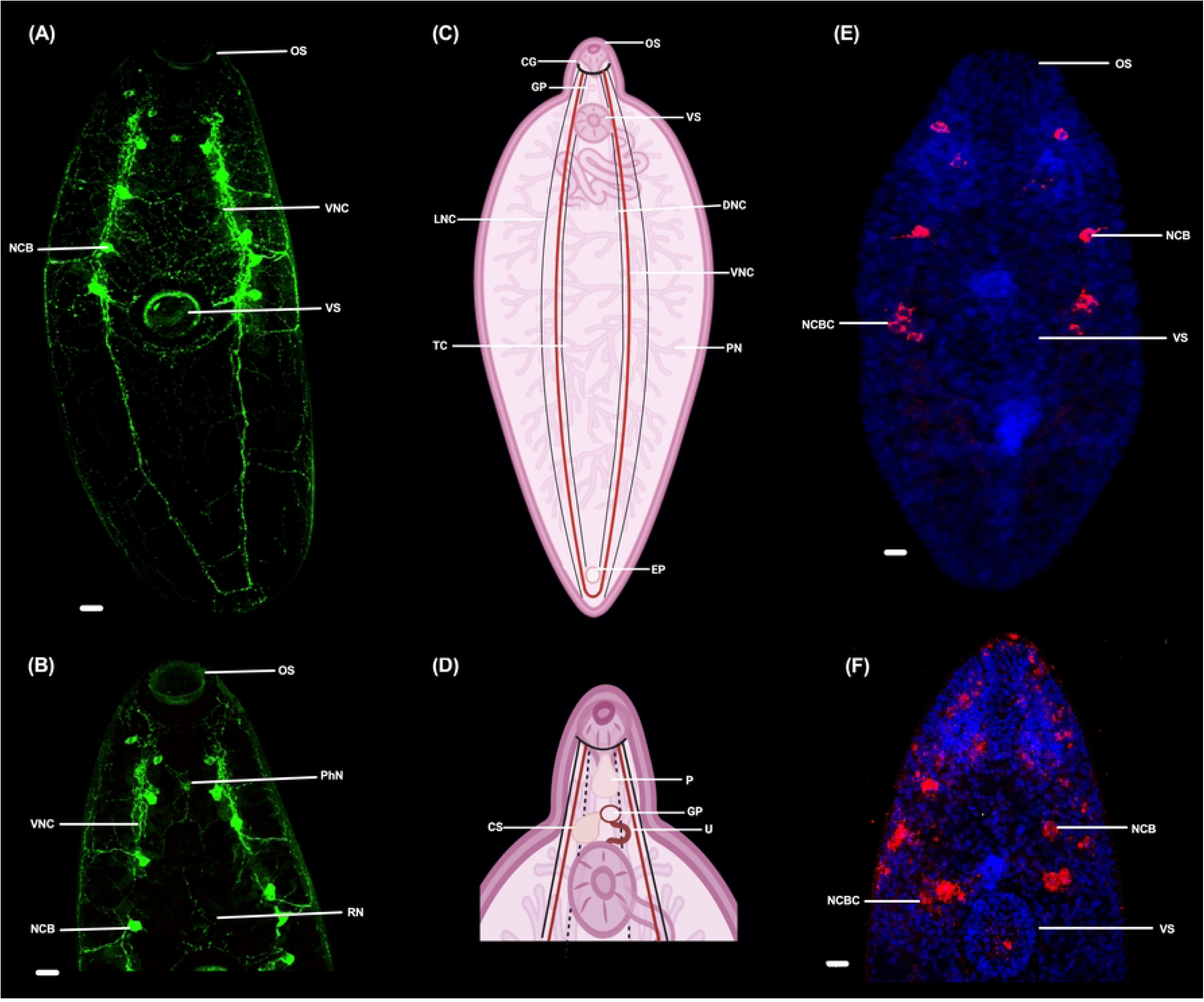
Serotonin and serotonin-signaling components are widely expressed in neurons in juvenile *Fasciola hepatica.* **(A) Whole-body immunocytochemistry in 21-day-old juvenile *F. hepatica* showing widespread serotonin immunoreactivity (IR), especially in the main ventral nerve cords and brain. Green fluorescence represents serotonin IR. (B) Anterior region immunolocalization of serotonin in 21-day-old juvenile *F. hepatica*, highlighting serotonin in the cerebral ganglia and surrounding neural structures.** Green fluorescence indicates serotonin IR. **(C) Whole-body schematic representation of *F. hepatica* adult, illustrating key biological structures and major nervous system components, including nerve cords, commissures, and neuronal cell bodies. (D) Anterior schematic of *F. hepatica,* showing essential oral and reproductive structures, including the oral sucker, pharynx, gonopore, and cirrus sac. (E) Whole-body fluorescent *in situ* hybridization (ISH) showing *F. hepatica* tryptophan hydroxylase (FhTPH) in the nervous system.** Red fluorescence indicates positive ISH signal, while blue fluorescence (DAPI staining) marks cell nuclei. FhTPH expression is observed in a pattern that aligns to neuronal cells of the nervous systems. **(F) *In situ* hybridization (ISH) showing expression of the *F. hepatica* serotonin-gated G protein-coupled receptor (5HT_1AFhep_) in anterior neurons and adjacent regions.** Red fluorescence shows receptor gene expression, while blue fluorescence (DAPI staining) highlights nuclei. Receptor localization is evident in the cerebral ganglia and other cells consistent with neuronal patterning. Abbreviations: **OS** = oral sucker, **VS** = ventral sucker, **VNC** = ventral nerve cord, **NCB** = neuronal cell body, **PhN** = pharyngeal nerves, **RN** = reproductive nerves, **CG** = cerebral ganglion, **GP** = gonopore, **DNC** = dorsal nerve cord, **LNC** = lateral nerve cord, **TC** = transverse commissure, **PN** = peripheral nerves, **P** = pharynx, **U** = uterus, **CS** = cirrus sac, **NCBC** = neuronal cell body cluster.

Fluorescent *in situ* hybridization (FISH) analysis demonstrated distinct spatial expression of FhTPH and 5HT_7CFhep_ GPCR transcripts within the nervous system of *F. hepatica* (Figure 4E and 4F). Transcript localization (Figure 4C and 4D) identified FhTPH expression predominantly in the ventral nerve cords and associated neuronal cell bodies (Figure 4E), consistent with neuronal-based serotonin biosynthesis (57). Similarly, 5-HT_7CFhep_ GPCR expression was localized to regions within the ventral nerve cords and adjacent areas peripheral to the cells expressing FhTPH transcripts, supporting the functional coupling of serotonin synthesis and signaling via GPCRs in these regions (Figure 4F).

Both transcripts localized predominantly to the anterior end of the parasite, particularly surrounding tissues critical for host interaction and reproduction, such as the pharynx, oral suckers, and reproductive organs (Figure 4C-F). Comparison of transcript distribution using *in situ* images indicated similar spatial expression patterns for FhTPH and 5HT_7CFhep_ GPCR transcripts in juvenile fluke signaling (S5 Figure).

### Elevated 5-HT levels enhance motility in juvenile *F. hepatica*

Exposure to 5-HT for 24 hours, in the absence of growth-enhancing media (RPMI), significantly increased the motility of *F. hepatica* juveniles by over 130% at a concentration of 1 mM (Figure 5A). This suggests that juvenile liver fluke can uptake exogenous 5-HT, a process that has also been characterized in other flatworm species (37,58). In *S. mansoni*, this ability has been attributed to the serotonin transporter SmSERT, which facilitates the uptake of 5-HT from the surrounding environment (58). While higher concentrations (>1 mM) also increased motility, a significant decrease in growth and survival was observed, indicating potential toxic effects at elevated levels (S6 Figure). Dose-dependent effects of 5-HT have been reported in other parasitic species, highlighting the delicate balance required for physiological relevance (30,48).

**Figure 5.**
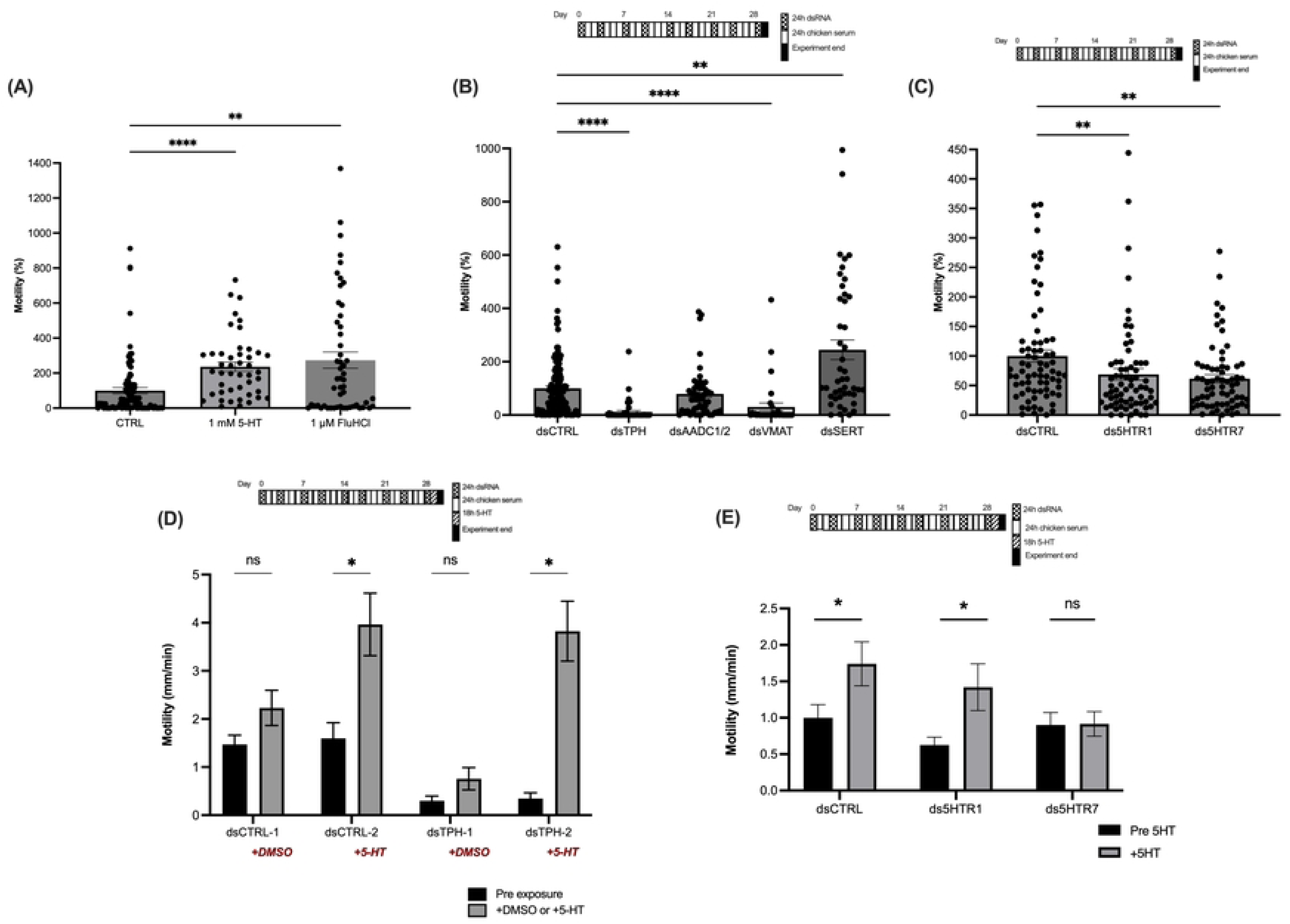
Chemical inhibition and gene-silencing of serotonin-signaling components in juvenile *Fasciola hepatica* alter motility. **(A) Increased motility in newly excysted juvenile *F. hepatica* following 24-hour exposure to 1 mM serotonin (5-HT) and 1 μM fluoxetine hydrochloride (FluHCL)**. Each data point represents motility (percentage length change per mm relative to the negative control) of an individual worm. Statistical significance determined by ANOVA; **p ≤ 0.01, ****p ≤ 0.0001. **(B) Altered motility of 28-day-old juvenile *F. hepatica* after RNAi knockdown of serotonin-signaling pathway genes.** Targeted genes include FhTPH (tryptophan hydroxylase), FhAADC1 and FhAADC2 (aromatic amino acid decarboxylase enzymes), FhVMAT (vesicular monoamine transporter), and FhSERT (serotonin reuptake transporter). Motility is expressed as percentage movement (mm/min) of individual juveniles compared to untreated controls. dsRNA exposure carried out twice per week as shown. Statistical significance determined by ANOVA; **p ≤ 0.01, ****p ≤ 0.0001. **(C) Decreased motility of 28-day-old juvenile *F. hepatica* following combinatorial RNAi knockdown of serotonin-G protein-coupled receptors (GPCRs).** Targeted receptors include type 1 (5HTR1AFhep and 5HTR1BFhep) and type 7 (5HTR7AFhep, 5HTR7BFhep, and 5HTR7CFhep) serotonin receptors. Data points represent motility (percentage, mm/min) of individual juveniles compared to untreated controls. dsRNA carried out twice per week as shown. Statistical significance determined by ANOVA; **p ≤ 0.01. **(D) Serotonin rescues the motility impact of FhTPH-knockdown in *F. hepatica* juveniles.** Black bars represent basal motility (mm/min) of juveniles post repeated double stranded (ds)RNA exposure (dsCTRL or dsTPH) but pre-chemical exposure. Gray bars show basal motility (mm/min) of juveniles post-dsRNA treatment followed by 18-hour exposure to 0.001% dimethyl sulfoxide (DMSO) or 1 mM 5-HT as outlined in red. Statistical significance determined by t-test; *p ≤ 0.05. **(E) The silencing of Fh5HT type 7 GPCRs abolishes the stimulatory effects of 1 mM 5-HT on juvenile liver fluke**. Black bars represent percentage motility (compared to RPMI) of juveniles post repeated dsRNA exposure (dsCTRL, ds5HTR1 [5HTR1AFhep, 5HTR1BFhep] or ds5HTR7 [5HTR7AFhep, 5HTR7BFhep, and 5HTR7Cfhep]) but pre-chemical exposure. Gray bars show motility after dsRNA treatment followed by 18-hour exposure to 1 mM 5-HT. Statistical significance determined by t-test; *p ≤ 0.05.

The motility-enhancing effects of 5-HT were further corroborated by blocking serotonin reuptake from synapses using fluoxetine hydrochloride (FluHCl), a selective serotonin reuptake inhibitor (SSRI). Following a 24-hour incubation with 1 μM FluHCl, juvenile motility increased by over 170% (Figure 5A). Importantly, lower concentrations of FluHCl also elicited significantly increased motility, suggesting high affinity of this compound for the serotonin transporter (FhSERT) in *F. hepatica* (S6 Figure). This is contrary to data showing low affinity binding of fluoxetine to SmSERT in *S. mansoni* (18).

*F. hepatica* juvenile motility was further enhanced by over 230% when culture media (50% CS) was supplemented with 1 mM 5-HT and maintained for 28 days (S7 Figure). This long-term exposure underscores the potential for sustained 5-HT availability to modulate neuromuscular activity in juvenile stages of *F. hepatica*. Together, these data suggest that serotonin enhances juvenile motility, consistent with a role in regulating neuromuscular functions in parasitic flatworms.

### Disruption of 5-HT signaling pathway components via RNAi induces aberrant motility in juvenile *F. hepatica*

All studied genes were successfully knocked down by more than or equal to 50%, as confirmed by transcript quantification (S8 Figure). The transcript reduction of key 5-HT signaling pathway components had profound effects on the motility of *F. hepatica* juveniles, underscoring the role of 5-HT in neuromuscular coordination. Knockdown of the 5-HT synthesis enzyme, FhTPH, resulted in a significant 84% reduction in juvenile motility, revealing the importance of serotonin biosynthesis in the maintenance of motor activity. Considering the phenotypic impacts of FhTPH-RNAi, it was surprising that silencing FhAADC1 and FhAADC2 did not significantly affect fluke motility, potentially reflecting that these enzymes are not rate limiting and/or there are unknown compensatory mechanisms/functional redundancy within endogenous dopaminergic/serotonergic systems; note that FhAADC enzymes also play roles in dopamine biosynthesis (15). As hypothesized, knockdown of the vesicular monoamine transporter, FhVMAT, responsible for serotonin storage and release, caused a significant 58.5% reduction in juvenile motility (Figure 5B). This supports the hypothesis that impaired VMAT function compromises neurotransmitter availability at the synapse. While TPH and VMAT have been identified in other parasites (15,43), this study provides the first characterization of their functional importance in a parasitic flatworm. In contrast, knockdown of the 5-HT reuptake transporter, FhSERT, likely to impede serotonin removal from synaptic junctions, led to a dramatic 235% increase in juvenile motility (Figure 5B), providing the first SERT-associated phenotype in a parasitic flatworm and confirming its importance in 5-HT signaling and homeostasis in *F. hepatica*.

To investigate their functional contributions to fluke motility, receptor clades were targeted using RNAi in a combinatorial approach: dsRNAs targeting both type 1 GPCRs (Fh5HT_1AFhep_ and Fh5HT_1BFhep_) were applied simultaneously, while a separate soak with three dsRNAs targeted the type 7 GPCRs (Fh5HT_7AFhep_, Fh5HT_7BFhep_, and Fh5HT_7CFhep_). Knockdown of the 5-HT type 1 receptors resulted in a significant 47.7% reduction in motility, while silencing the three type 7 receptors led to an even more pronounced 60% reduction in motility (Figure 4C). These findings support the hypothesis that 5-HT GPCRs play a role in modulating neuromuscular activity in juvenile *F. hepatica*.

### Exogenous 5-HT restores motility in FhTPH RNAi-treated parasites and validates 5-HT GPCR function

Knockdown of FhTPH significantly reduces the motility of juvenile *F. hepatica*, indicating an impaired ability to synthesize 5-HT endogenously. Supporting this hypothesis, the exogenous application of 1 mM 5-HT for 18 hours restores and enhances the motility of FhTPH knockdown juveniles by tenfold, supporting the RNAi phenotype data (Figure 5D). Similar methods were employed to validate the functional roles of 5HT receptors. Exogenous application of 5-HT increased motility and partially rescued the motility deficits of juveniles with silenced 5HT_1_ receptors (Figure 5E). A significant 50% enhancement was observed, suggesting that 5-HT-mediated motility is primarily associated with the activity of 5HT_7_ receptors. This hypothesis was further supported by the lack of motility response to exogenous 5-HT in juveniles with silenced 5HT_7_ receptors (Figure 5E). These findings align with the functional characterization of the *S. mansoni* 5-HT₇ receptor, where RNAi-mediated silencing resulted in a marked reduction in schistosomula motility, underscoring the receptor’s critical role in movement regulation (30). Similarly, RNAi knockdown of 5HT₁ receptors revealed a significant but less pronounced impact on motility in *F. hepatica* juveniles (Figure 5C). These data collectively suggest that compared to 5HT₁ receptors, the 5HT_7_ receptors play a more significant role in the direct modulation of juvenile motility, providing the first functional delineation of 5-HT receptor subtypes in parasitic flatworms. These observations are consistent with vertebrate studies, where 5HT₁ receptors are characterized as ’autoregulatory’ receptors (59). In vertebrates, they are integral to the homeostatic regulation of serotonin signaling through coupling with Gi/o inhibitory proteins, highlighting their indirect but essential role in neurotransmission (52,59).

### Elevated 5-HT levels enhance growth of juvenile *F. hepatica*

Interestingly, juveniles maintained in a medium consisting of 50% chicken serum supplemented with 1 mM 5-HT over a 28-day period exhibited a significant 30.8% increase in growth (Figure 6A and 6B). This finding suggests that elevated 5-HT levels can enhance the growth of juvenile *F. hepatica*. 5-HT is well-documented as a critical neuromodulator in invertebrates, with roles extending beyond neurotransmission to include the regulation of growth, reproduction, and metabolism. In parasitic flatworms, previous studies have highlighted serotonin’s role in modulating motility and host-parasite interactions, but its influence on growth has been less explored. There are a few studies that suggest 5-HT may influence cell proliferation and tissue growth through its influence on energy metabolism and cell division (32,33,35,36). 5-HT stimulates glucose uptake and promotes glycogen breakdown in *F. hepatica* and *S. mansoni*, leading to increased lactic acid production under anaerobic conditions (60,61). This metabolic shift suggests that 5-HT enhances glycolysis, ensuring a rapid energy supply necessary for energy-intensive processes such as motility. The resulting increase in metabolic activity may also support the growth and development of these flatworms. Emerging research suggests that 5-HT plays a pivotal role in directly regulating cell proliferative mechanisms across a range of species. Although evidence remains limited in invertebrates, studies in *E. multilocularis* have demonstrated that 5-HT signaling is essential for larval development and promotes the expansion of germinal layer cells, which are critical for parasite growth and survival (37). In vertebrates, the role of 5-HT in cell proliferation is more extensively documented, particularly in the context of cancer – for recent review see (62). For example, in the central nervous system, 5-HT modulates neurogenesis by regulating both the proliferation and differentiation of neural progenitor cells (63,64), whilst treating non-small cell lung cancer cells with 5-HT enhanced their proliferation and migration (65). Here, the growth-enhancing effect of 5-HT was substantiated for *F. hepatica,* as the growth of juveniles increased significantly by 29.1% following RNAi-mediated knockdown of the 5-HT transporter gene (FhSERT) (Figure 6C). No significant change in growth was observed with knockdown of other 5-HT pathway genes, highlighting the importance of 5-HT transport in modulating serotonergic signaling and its downstream effects on parasite biology. RNAi-mediated knockdown of GPCRs provides further evidence of the role of 5-HT in developmental processes, particularly through its interaction with 5HT_1_ receptors. A significant 22.3% increase in juvenile size was observed when 5HT_1_ receptors were silenced (Figure 6D), suggesting 5HT_1_ receptors may function as inhibitory regulators of growth, where their silencing removes this inhibition, thus enhancing growth. While the precise pathways of action remain under investigation, existing research suggests that, beyond its well-documented role in inhibiting cAMP production, 5-HT is directly involved in regulating multiple cell cycle progression pathways (62). Notably, it has been linked to the mitogen-activated protein kinase (MAPK) and PI3K/Akt signaling cascades, which are essential for cellular growth, survival and differentiation (66), as well as the Hippo signaling pathway, a key regulator of organ size and cell proliferation (67). These interactions are thought to be mediated by specific 5-HT receptor subtypes, particularly 5HT_7_ and 5HT_1_ receptors (68,69).

The interplay between serotonin receptor subtypes in regulating developmental processes in fluke underscores the complexity of 5-HT signaling. Future research should aim to elucidate the precise molecular mechanisms underlying 5-HT-mediated growth in juvenile liver fluke, with a particular focus on identifying downstream pathways involved in cell proliferation and tissue differentiation. A deeper understanding of these processes could significantly expand the repertoire of drug targets associated with serotonin signaling.

**Figure 6.**
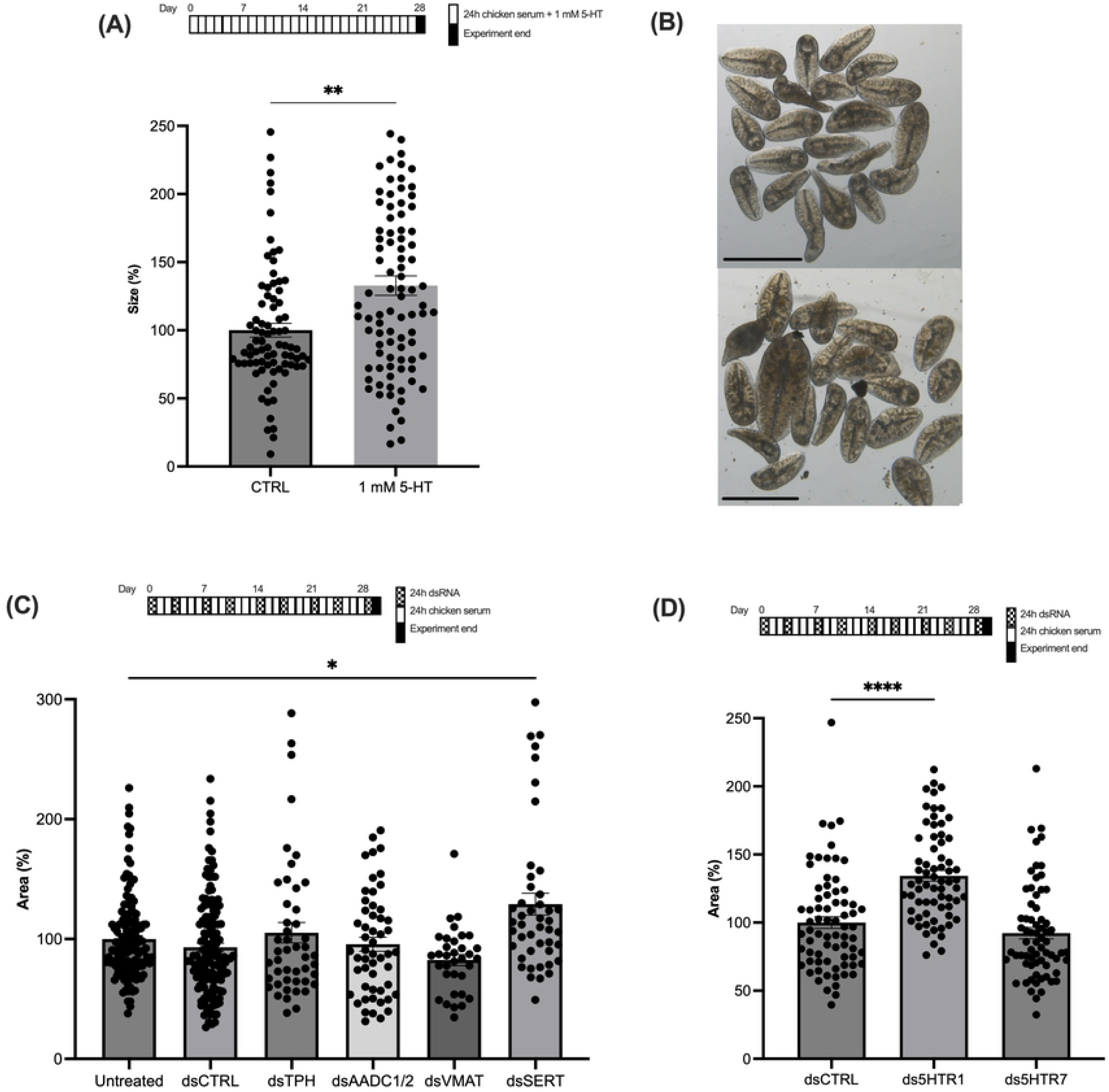
Serotonin and the gene silencing of selected serotonin-signaling components alter juvenile *Fasciola hepatica* growth. **(A) The growth enhancing effect of prolonged exposure (28 days) to 1 mM serotonin (5-HT) on juvenile *F. hepatica*.** Each data point represents the size of an individual juvenile expressed as percentage area (mm²) relative to the unsupplemented media control. Statistical significance determined by t-test analysis; **p ≤ 0.01. **(B) Brightfield images showing the growth enhancing impact of 1 mM 5-HT supplementation on 21-day-old *F. hepatica* juveniles**. Top image shows juveniles without 5-HT supplementation, bottom image shows juveniles with 5-HT supplementation. **(C) Growth enhancing effect of FhSERT silencing in 28-day-old *F. hepatica* juveniles.** Targeted genes include FhTPH (tryptophan hydroxylase), FhAADC1 and FhAADC2 (aromatic amino acid decarboxylases), FhVMAT (vesicular monoamine transporter), and FhSERT (serotonin reuptake transporter). Growth is expressed as percentage area (mm²) of individual juveniles compared to untreated controls. dsRNA exposure carried out twice a week as shown. Statistical significance determined by ANOVA; *p ≤ 0.05. **(D) RNAi-mediated knockdown of type 1 serotonin-G protein-coupled receptors (GPCRs) in 28-day-old juvenile *F. hepatica* enhanced growth.** Targeted receptors include type 1 (5HTR1AFhep and 5HTR1BFhep) and type 7 (5HTR7AFhep, 5HTR7BFhep, and 5HTR7CFhep). Each data point represents the relative area (percentage, mm²) of an individual juvenile compared to untreated controls. dsRNA exposure carried out twice a week as shown. Statistical significance determined by ANOVA; ****p ≤ 0.0001.

### Conclusion

The findings of this study provide compelling evidence that serotonin (5-HT) plays a pivotal role in regulating not only neuromuscular function, but also growth and development in *F. hepatica*. These findings establish serotonin as a fundamental regulator of *F. hepatica* biology, offering new insights into its potential as a therapeutic target for parasite control.

## Methods

### Computational characterization of serotonergic signaling pathway of *F. hepatica*

Key genes involved in serotonergic signaling were identified using BLAST analyses of the *Fasciola hepatica* predicted protein datasets (PRJEB58756) with characterized human (TPH; P17752, AADC; P20711, VMAT; Q05940, SERT P31645) and *Schistosoma mansoni* (TPH; Smp_174920, AADC; Smp_171580, VMAT; Smp_121920, SERT; Smp_333840) orthologs as queries (15,15,18,39–43,70). Predicted protein datasets used in this study were downloaded from WormBase ParaSite (version WBPS19; https://parasite.wormbase.org/index.html) (71).

Annotated GPCR sequences (15,30,44,72) were used to interrogate the genomes of *F. hepatica* and 28 other flatworm species (16 cestodes; *Echinococcus canadensis* (PRJEB8992), *Echinococcus granulosus* (PRJNA182977 - G1), *Echinococcus multilocularis* (PRJEB122 - Java), *Echinococcus oligarthrus* (PRJEB31222 - DaMi1), *Hymenolepis diminuta* (PRJEB507 - Denmark), *Hymenolepis microstoma* (PRJEB124), *Hymenolepis nana* (PRJEB508 - Japan), *Mesocestoides corti* (PRJEB510 - Specht & Voge 1965), *Taenia asiatica* (PRJEB532 - South Korea), *Taenia multiceps* (PRJNA307624 - Gns01), *Taenia saginata* (PRJNA71493 - TSAYD01), *Taenia solium* (PRJNA170813 - Mexico), *Dibothriocephalus latus* (PRJEB1206), *Spirometra erinaceieuropaei* (PRJEB1202), *Schistocephalus solidus* (PRJEB527 - NST_G2), *Hydatigera taeniaeformis* (PRJEB534 - Spain/Canary Islands) and 12 trematode species; *Clonorchis sinensis* (PRJNA386618 - Cs-k2), *Schistosoma bovis* (PRJNA451066), *Schistosoma japonicum* (PRJNA520774 - HuSjv2), *Fasciolopsis buski* (PRJNA284521 - HT), *Fasciola gigantica* (PRJNA230515 - Uganda_cow_1), *Schistosoma haematobium* (PRJNA78265), *Schistosoma curassoni* (PRJEB519 - Senegal/Dakar), *Schistosoma mansoni* (PRJEA36577), *Trichobilharzia regenti* (PRJEB44434 - tdTriRege1.1), *Opisthorchis viverrini* (PRJNA222628), *Paragonimus westermani* (PRJNA454344), *Echinostoma caproni* (PRJEB1207 - Egypt)) using Hidden Markov Models (HMM) to identify additional 5-HT receptor candidates (73). Protein hits with E-values <0.01 were validated through reciprocal BLAST searches against National Center for Biotechnology Information (NCBI) non-redundant (nr) database sans Platyhelminthes and UniProt datasets (70,74). GPCRs with ≥4 transmembrane domains were confirmed using TMHMM-2.0 (73), InterProScan (version 5; (75)), and conserved motif analysis. Expression levels were quantified as transcripts per million (TPM) from published datasets (46,71). Alignments were performed with Clustal Omega (76), and maximum likelihood phylogenetic trees were generated using MEGA (version 11; (77)) (model=WAG+F; bootstrap=500) and formatted using iTOL (78). Protein relationships were further visualized through CLANs analysis (79). GPCR structures were predicted using Phyre2.2 software (80) and imported into PyMOL (81) for visualization and ligand binding analysis using the PyMOL plugin DockingPie 1.2 (82). The serotonin (SRO) ligand structure was downloaded from Protein Data Bank (PDB) and imported directly into PyMOL for docking analysis (83)

### *F. hepatica* and excystment

Juvenile *F. hepatica* of Italian strain (obtained from Ridgeway Research) were prepared and maintained as follows: metacercariae were stored in distilled water at 4°C until needed and excysted following the protocol described by McVeigh et al. (84). Prior to excystment, metacercariae were manually popped from their outer wall casing and bleached in 10% sodium hypochlorite solution. Newly excysted juveniles (NEJs) were cultured at 37°C in a humidified atmosphere with 5% CO₂, either in 50:50 chicken serum:RPMI 1640 medium (CS: Merck, C5405; RPMI 1640; Thermo Fisher Scientific) or in RPMI 1640 alone, depending on the experimental application, as outlined by McCusker et al. (85). The culture medium was replaced three times per week during extended culture periods. The complete protocol can be accessed at: https://dx.doi.org/10.17504/protocols.io.14egn212qg5d/v1

### Immunocytochemistry

21 day old juveniles were flat fixed under a coverslip for 10 minutes at room temperature and free fixed overnight at 4 °C in 4% paraformaldehyde. Fixed juveniles were washed and stored in antibody diluent (AbD; 0.1 M phosphate-buffered saline (PBS) (Merck), 0.1 % Triton-X (Merck), 0.1 % bovine serum albumin (Merck)) at 4 °C until required. Serotonin primary antibody (S5545, Merck) incubations were carried out at 1:1000 dilution in AbD for 72 hours rotating at 4 °C. Juveniles were washed in AbD before incubation in fluorescein isothiocyanate (FITC)-labelled anti-rabbit secondary antiserum (Merck) at 1:100 concentration for 48 hours rotating at 4 °C. Juveniles were further washed in AbD before mounting in Vectashield (Vector Laboratories) and imaged on a Leica TCS SP8 confocal scanning laser microscope.

### Whole mount fluorescence *in situ* hybridization (FISH)

*In situ* hybridization methods were carried out as described in detail by Armstrong et al. (86) In summary, after 4 weeks of culture, juveniles were incubated with 500 µM 5-ethynyl-2′-deoxyuridine (EdU) in CS50 medium for 24 hours to label EdU+ cells for FISH co-localization. RNase-free pipette tips and DEPC-treated water were used up to the Riboprobe incubation step. Juveniles were fixed under a coverslip in 4% formaldehyde for 10 minutes, followed by free fixation on a rotator for another 10 minutes at room temperature. They were then washed in PBSTx (1× PBS with 0.3% Triton-X) and dehydrated in 50% methanol:PBSTx before storage in 100% methanol at −20°C. RNA probes for FhTPH and 5HT_1CFhep_ were synthesized from T7 templates amplified via PCR using the Roche FastStart kit. Probe primers are outlined in S4 Table. Probe synthesis involved overnight incubation at 27°C with DIG-UTP labelling mix (Roche), transcription buffer, and T7 RNA polymerase (T7 RNA polymerase kit, ThermoFisher Scientific), followed by DNase treatment (0.02%) and ethanol precipitation with 0.1 M lithium chloride at −80°C. RNA pellets were centrifuged (x 16000*g*, 4 °C), washed in 70% ethanol, resuspended in nuclease-free water, and assessed for purity and concentration using a DeNovix DS-11 FX spectrophotometer. Probes were diluted to 50 ng/µL in hybridization buffer (50% de-ionized formamide, 25% saline-sodium citrate buffer (SSC), 0.1 mg/ml yeast RNA, 1% (v/v) Tween-20, 5% dextran sulfate, made to volume with DEPC-treated H_2_O) and stored at -20 °C. Yeast RNA was prepared per Jing (87).

*In situ* hybridization methods were adapted from methods of *Schistosoma mansoni* and *Schmidtea mediterranea* (88). A full protocol is available at https://www.protocols.io/view/fluorescent-in-situ-hybridisation-for-juvenile-fas-3byl4q7xjvo5/v1. Flat-fixed juveniles were rehydrated with 10-minute washes in 50% PBSTx:MeOH and 1× SSC, then bleached (8.85 ml H_2_O, 0.5 ml formamide, 0.25 ml 20x SSC, 0.4 ml 30 % H_2_O_2_) under light for 1.5 hours. Worms were further washed for 10 minutes in 1x SSC and PBSTx. Juveniles were permeabilized with proteinase K solution (0.1% SDS and 0.01 mg/ml proteinase K) and post-fixed for 10 minutes in 4% formaldehyde. Worms were prepared into Intavis baskets (35 μm mesh) and incubated for 10 minutes in PBST:Pre-hybridization solution (50% de-ionized formamide, 25% SSC, 0.1 mg/ml yeast RNA, 1% (v/v) Tween-20, made up to volume with DEPC-treated H_2_O) before a 2-hour incubation at 52 °C in pre-hybridization solution. Probes were pre-heated at 80°C for 5 minutes and hybridized with juveniles overnight (≥16 h) at 52°C using 1 ng/µl probe solution; sense probes served as controls. Post-hybridization, worms were washed in 50% hybridization solution: 2x SSCx (2x SSC with 0.1% Triton X-100), 2x SSCx and 0.2x SSCx at 52 °C, followed by two 10-minute TNTx (0.1 M Tris pH 7.5, 0.15 M NaCl, 0.1% Tween-20) washes at room temperature. Blocking was performed with 5% inactivated horse serum and 0.5% Western Blocking Reagent in TNTx for 2 hours, followed by overnight incubation at 4°C with 1:2000 anti-DIG-POD (diluted in blocking solution, Merck). Worms underwent sequential TNTx washes (5, 10 and 6x 20 minutes) at room temperature and were incubated in tyramide solution (TSA buffer; 2 M NaCl; 0.1 M Boric acid pH 8.5, 0.5% H_2_O_2_, 0.1% 4-Iodophenylboronic acid (4IPBA) 20 mg/ml, 1:500 5-TAMRA (5-carboxytetramethylrhodamine, Merck) for 10 minutes at room temperature. Worms were stained with 4,6-diamidino-2-phenylindole (DAPI) overnight and washed in TNTx for 48 hours. Juveniles were mounted in Vectashield before imaging using a Leica TCS SP8 confocal microscope.

### Chemical exposures

Serotonin hydrochloride powder (H9523, Merck) was dissolved to 100 mM stock concentration in milliQ water. For short term exposures, juveniles were exposed to a final concentration of 0.01 μM – 1 mM serotonin in 3 ml RPMI. For long exposures, juveniles were cultured in 200 μl 50% CS supplemented with 0.01 μM – 1 mM serotonin. Fluoxetine hydrochloride was dissolved to 10 mM stock concentration in dimethyl sulfoxide (DMSO) and further diluted to desired final concentrations (0.01 μM – 1 mM) in 3 ml RPMI 1640. 0.02% ddH_2_O and 0.001% DMSO were used as negative controls as required. Images and videos were captured using an Olympus SZX10 stereo microscope with CAM-SC50 camera to monitor phenotypic changes (area-mm^2^; motility-length change, mm/min) determined via ImageJ analysis using WrMTrcK plugin (89,90).

### RNA interference

*F. hepatica* juvenile RNAi was carried out as described by McCusker et al. (91). Double stranded RNAs (dsRNA) specific to bacterial neomycin (bacterial neomycin phosphotransferase, U55762), red fluorescent protein (cloning vector pAJ50, ARW80046), *F. hepatica* tryptophan hydroxylase (FhTPH; FhHiC23_g11603), *F. hepatica* aromatic decarboxylase 1&2 (FhAADC1; FhHiC23_g50, FhAADC2; FhHiC23_g2511), *F. hepatica* vesicular monoamine transporter (FhVMAT; FhHiC23_g2), *F. hepatica* serotonin transporter (FhSERT; FhHiC23_g1585) and five serotonin receptors (FhHiC23_g4917, FhHiC23_g1196, FhHiC23_g115, FhHiC23_g2340, FhHiC23_g7880) were generated using the T7 RiboMAX™ Express RNAi System (Promega) from cDNA templates labelled with the T7 promoter sequence; 5’-TAATACGACTCACTATAGGGT-3’ amplified by PCR. All primers are listed in S4 Table. All RNAi experiments were carried out at least in triplicate. Juveniles were exposed to 100 ng/μl dsRNA (target or control) in RPMI 1640 for 24 hours twice per week for 4 weeks under normal culture conditions. Between dsRNA exposures juveniles were cultured as standard in 50% CS. Post RNAi exposures, juveniles were snap frozen for extraction and transcript quantification.

Phenotypic analysis was carried out at day 29 post excystment after dsRNA exposure, comparing to RPMI and dsRNA control groups. Juveniles were snap frozen in liquid nitrogen for transcript analysis.

### Transcript analysis

mRNA was extracted from RNAi treatment groups using Dynabeads mRNA direct Kit (Thermo Fisher Scientific). cDNA was synthesized (High Capacity RNA-to-cDNA kit, Thermo Fisher Scientific) following DNase treatment (Turbo DNA-free, Thermo Fisher Scientific). qPCRs were performed on a Rotor-Gene Q 5-plex HRM PCR system (Qiagen) using SensiFast SYBR (Bioline) and primers outlined in S4 Table at final concentration 1.25 μM. Cycling parameters; 10 minutes at 95 °C, followed by 40 cycles of 95 °C 10 s, 60 °C 15 s, 72 °C 30 s. All PCRs were performed in triplicate and included no-template controls and melt-curve analyses as standard. Relative expression analysis was determined using Pfaffl’s Augmented ΔΔCt method (92), normalizing expression in each sample relative to the untreated control, standardized to a glyceraldehyde 3-phosphate dehydrogenase (GAPDH, FhHiC23_g5828) housekeeping gene. Statistical significance was determined relative to the effects of negative control treatments (dsCTRL) on target gene expression.

### Statistical Analysis

Growth and motility data were analyzed using one-way ANOVA with Dunnett’s post hoc test, analyzing significance of treatments relative to the negative control dsRNA treated juveniles (RNAi experiments), or vehicle control treated juveniles (compound exposure experiments). All raw data for figures can be found in S3 Table.

## Supplementary

**S1 Figure. Alignment of flatworm serotonin (5-HT) gated G protein coupled receptors S1 Table. Summary of bioinformatic analysis defining serotonin signaling genes in *Fasciola hepatica***

**S2 Figure. Motif analysis of *Fasciola hepatica* serotonin receptors**

**S2 Table. Flatworm serotonin (5-HT) gated G protein coupled receptor gene sequences S3 Figure. G protein coupling predictions for *Fasciola hepatica* serotonin G protein coupled receptors**

**S3 Table. Raw data for manuscript Figures 5 and 6**

**S4 Figure. PyMOL alignment model of human serotonin receptors and Fasciola hepatica serotonin receptors**

**S4 Table. Primer details for amplification of *Fasciola hepatica* genes targets for RNA interference, qPCR analysis and in situ hybridisation**

**S5 Figure. Neuronal mapping of cells expressing genes associated with serotonin signaling in *Fasciola hepatica* with fluorescent in situ hybridization**

**S6 Figure. Optimization of serotonin (5-HT) exposure for *Fasciola hepatica* juveniles S7 Figure. Motility of Fasciola hepatica juveniles after long term exposure to serotonin (5-HT)**

**S8 Figure. Knockdown values (ΔΔCt) for *Fasciola hepatica* serotonin signaling genes post RNA interference**

## References

1. Food and Agriculture Organisation of the United Nations. Diseases in Domestic Animals Caused by Flukes. Rome: Food and Agriculture Organisation. 1994.

2. Tolan RW. Fascioliasis due to *Fasciola hepatica* and *Fasciola gigantica* infection: an update on this Neglected Tropical Disease. Lab Med. 2011 Feb;42(2):107–16.

3. González-Miguel J, Becerro-Recio D, Siles-Lucas M. Insights into *Fasciola hepatica* juveniles: crossing the Fasciolosis rubicon. Trends Parasitol. 2021 Jan;37(1):35–47.

4. Kelley JM, Elliott TP, Beddoe T, Anderson G, Skuce P, Spithill TW. Current threat of triclabendazole resistance in *Fasciola hepatica*. Trends Parasitol. 2016 Jun;32(6):458– 69.

5. Beesley NJ, Cwiklinski K, Allen K, Hoyle RC, Spithill TW, La Course EJ, et al. A major locus confers triclabendazole resistance in *Fasciola hepatica* and shows dominant inheritance. PLoS Pathog. 2023 Jan 26;19(1):e1011081.

6. McVeigh P, McCusker P, Robb E, Wells D, Gardiner E, Mousley A, et al. Reasons to be nervous about flukicide discovery. Trends Parasitol. 2018 Mar;34(3):184–96.

7. McVeigh P, Atkinson L, Marks NJ, Mousley A, Dalzell JJ, Sluder A, et al. Parasite neuropeptide biology: seeding rational drug target selection? Int J Parasitol Drugs Drug Resist. 2012 Dec;2:76–91.

8. Wolstenholme AJ. Ion channels and receptor as targets for the control of parasitic nematodes. Vol. 1, International Journal for Parasitology: Drugs and Drug Resistance. 2011. p. 2–13.

9. Geary TG, Klein RD, Vanover L, Bowman JW, Thompson DP. The nervous systems of helminths as targets for drugs. J Parasitol. 1992 Apr;78(2):215.

10. Okaty BW, Commons KG, Dymecki SM. Embracing diversity in the 5-HT neuronal system. Nat Rev Neurosci. 2019 Jul 4;20(7):397–424.

11. Jones LA, Sun EW, Martin AM, Keating DJ. The ever-changing roles of serotonin. Int J Biochem Cell Biol. 2020 Aug;125:105776.

12. Frazer A, Hensler J. Serotonin. In: Siegel GJ, Agranoff BW, Albers WR, Fisher SK, Uhler MD, editors. Basic Neurochemistry, Molecular, Cellular and Medical Aspects. 6th Edition. Philadelphia: American Society for Neurochemistry; 1999.

13. Millan M, Marin P, Bockaert J, Mannourylacour C. Signaling at G-protein-coupled serotonin receptors: recent advances and future research directions. Trends Pharmacol Sci. 2008 Sep;29(9):454–64.

14. Ribeiro P, El-shehabi F, Patocka N. Classical transmitters and their receptors in flatworms. Parasitology. 2006 Mar 29;131(S1):S19.

15. Preza M, Montagne J, Costábile A, Iriarte A, Castillo E, Koziol U. Analysis of classical neurotransmitter markers in tapeworms: evidence for extensive loss of neurotransmitter pathways. Int J Parasitol. 2018 Nov;48(13).

16. Bennett JL, Bueding E. Uptake of 5-hydroxytryptamine by *Schistosoma mansoni*. Mol Pharmacol. 1973 May;9(3):311–9.

17. Catto BA, Ottesen EA. Serotonin uptake in schistosomules of *Schistosoma mansoni*. Comparative Biochemistry and Physiology Part C: Comparative Pharmacology. 1979 Jan;63(2):235–42.

18. Patocka N, Ribeiro P. Characterization of a serotonin transporter in the parasitic flatworm, *Schistosoma mansoni*: cloning, expression and functional analysis. Mol Biochem Parasitol. 2007 Aug;154(2):125–33.

19. Fairweather I, Maule AG, Mitchell SH, Johnston CF, Halton DW. Immunocytochemical demonstration of 5-hydroxytryptamine (serotonin) in the nervous system of the liver fluke, *Fasciola hepatica* (Trematoda, Digenea). Parasitol Res. 1987;73(3):255–8.

20. Sukhdeo SC, Sukhdeo MVK. Immunohistochemical and electrochemical detection of serotonin in the nervous system of *Fasciola hepatica*, a parasitic flatworm. Brain Res. 1988 Oct;463(1):57–62.

21. Tembe EA, Holden-Dye L, Smith SWG, Jacques PAM, Walker RJ. Pharmacological profile of the 5-hydroxytryptamine receptor of *Fasciola hepatica* body wall muscle. Parasitology. 1993 Jan 6;106(1):67–73.

22. Holmes SD, Fairweather I. *Fasciola hepatica*: The effects of neuropharmacological agents upon in vitro motility. Exp Parasitol. 1984 Oct;58(2):194–208.

23. Thompson CS, Mettrick DF. The effects of 5-hydroxytryptamine and glutamate on muscle contraction in *Hymenolepis diminuta* (Cestoda). Can J Zool. 1989 May 1;67(5):1257–62.

24. Maule AG, Halton DW, Allen JM, Fairweather I. Studies on motility *in vitro* of an ectoparasitic monogenean, *Diclidophora merlangi*. Parasitology. 1989 Feb 6;98(1):85– 93.

25. Boyle JP, Zaide J V., Yoshino TP. *Schistosoma mansoni*: effects of serotonin and serotonin receptor antagonists on motility and length of primary sporocysts *in vitro*. Exp Parasitol. 2000 Apr;94(4):217–26.

26. McKay DM, Halton DW, Allen JM, Fairweather I. The effects of cholinergic and serotoninergic drugs on motility *in vitro* of *Haplometra cylindracea* (Trematoda: Digenea). Parasitology. 1989 Oct 6;99(2):241–52.

27. Hrčkova G, Velenbný S, Halton DW, Maule AG. *Mesocestoides corti* (syn. *M. vogae*): modulation of larval motility by neuropeptides, serotonin and acetylcholine. Parasitology. 2002 Apr 1;124(4):409–21.

28. Pax RA, Siefker C, Bennett JL. *Schistosoma mansoni*: differences in acetylcholine, dopamine, and serotonin control of circular and longitudinal parasite muscles. Exp Parasitol. 1984 Dec;58(3):314–24.

29. Day TA, Bennett JL, Pax RA. Serotonin and its requirement for maintenance of contractility in muscle fibres isolated from *Schistosoma mansoni*. Parasitology. 1994 May 6;108(4):425–32.

30. Patocka N, Sharma N, Rashid M, Ribeiro P. Serotonin signaling in *Schistosoma mansoni*: a serotonin–activated G protein-coupled receptor controls parasite movement. PLoS Pathog. 2014 Jan 16;10(1):e1003878.

31. Rahman MS, Mettrick DF, Podesta RB. Effects of 5-hydroxytryptamine on carbohydrate metabolism in *Hymenolepis diminuta* (Cestoda). Can J Physiol Pharmacol. 1983 Feb 1;61(2):137–43.

32. Mansour TE. The effect of serotonin and related compounds on the carbohydrate metabolism of the liver fluke, *Fasciola hepatica*. J Pharmacol Exp Ther. 1959 Jul;126(3):212–6.

33. Rahman MS, Mettrick DF, Podesta RB. *Schistosoma mansoni*: effects of *in vitro* serotonin (5-HT) on aerobic and anaerobic carbohydrate metabolism. Exp Parasitol. 1985 Aug;60(1):10–7.

34. Karki S, Saadaoui M, Dunsing V, Kerridge S, Da Silva E, Philippe JM, et al. Serotonin signaling regulates actomyosin contractility during morphogenesis in evolutionarily divergent lineages. Nat Commun. 2023 Sep 8;14(1):5547.

35. Balakrishna P, George S, Hatoum H, Mukherjee S. Serotonin pathway in cancer. Vol. 22, International Journal of Molecular Sciences. MDPI AG; 2021. p. 1–10.

36. Sarkar A, Mukundan N, Sowndarya S, Dubey VK, Babu R, Lakshmanan V, et al. Serotonin is essential for eye regeneration in planaria *Schmidtea mediterranea*. FEBS Lett. 2019 Nov 27;593(22):3198–209.

37. Herz M, Brehm K. Serotonin stimulates *Echinococcus multilocularis* larval development. Parasit Vectors. 2021 Dec 6;14(1):14.

38. Marchant JS, Harding WW, Chan JD. Structure-activity profiling of alkaloid natural product pharmacophores against a Schistosoma serotonin receptor. Int J Parasitol Drugs Drug Resist. 2018 Dec;8(3):550–8.

39. Boularand S, Darmon MC, Ganem Y, Launay JM, Mallet J. Complete coding sequence of human tryptophan hydroxylase. Nucleic Acids Res. 1990;18(14):4257–4257.

40. Ichinose H, Kurosawa Y, Titani K, Fujita K, Nagatsu T. Isolation and characterization of a cDNA clone encoding human aromatic L-amino acid decarboxylase. Biochem Biophys Res Commun. 1989 Nov;164(3):1024–30.

41. Surratt CK, Persico AM, Yang XD, Edgar SR, Bird GS, Hawkins AL, et al. A human synaptic vesicle monoamine transporter cDNA predicts posttranslational modifications, reveals chromosome 10 gene localization and identifies *Taq* I RFLPs. FEBS Lett. 1993 Mar 8;318(3):325–30.

42. Lesch KP, Wolozin BL, Estler HC, Murphy DL, Riederer P. Isolation of a cDNA encoding the human brain serotonin transporter. J Neural Transm. 1993 Feb;91(1):67–72.

43. Hamdan FF, Ribeiro P. Characterization of a stable form of tryptophan hydroxylase from the human parasite *Schistosoma mansoni*. Journal of Biological Chemistry. 1999 Jul;274(31):21746–54.

44. McVeigh P, McCammick E, McCusker P, Wells D, Hodgkinson J, Paterson S, et al. Profiling G protein-coupled receptors of *Fasciola hepatica* identifies orphan rhodopsins unique to phylum Platyhelminthes. Int J Parasitol Drugs Drug Resist. 2018 Apr;8(1):87– 103.

45. McCorvy JD, Roth BL. Structure and function of serotonin G protein-coupled receptors. Pharmacol Ther. 2015 Jun;150:129–42.

46. Cwiklinski K, Dalton JP, Dufresne PJ, La Course J, Williams DJ, Hodgkinson J, et al. The *Fasciola hepatica* genome: gene duplication and polymorphism reveals adaptation to the host environment and the capacity for rapid evolution. Genome Biol. 2015 Dec 3;16(1):71.

47. Wheeler NJ, Heimark ZW, Airs PM, Mann A, Bartholomay LC, Zamanian M. Genetic and functional diversification of chemosensory pathway receptors in mosquito-borne filarial nematodes. PLoS Biol. 2020 Jun 1;18(6).

48. Camicia F, Celentano AM, Johns ME, Chan JD, Maldonado L, Vaca H, et al. Unique pharmacological properties of serotoninergic G-protein coupled receptors from cestodes. PLoS Negl Trop Dis. 2018 Feb 9;12(2).

49. Tierney AJ. Structure and function of invertebrate 5-HT receptors: a review. Vol. 128, Comparative Biochemistry and Physiology Part A. 2001.

50. Tierney AJ. Invertebrate serotonin receptors: A molecular perspective on classification and pharmacology. Vol. 221, Journal of Experimental Biology. Company of Biologists Ltd; 2018.

51. Kreshchenko N, Terenina N, Ermakov A. Serotonin signalling in flatworms: an immunocytochemical localisation of 5-HT7 type of serotonin receptors in *Opisthorchis felineus* and *Hymenolepis diminuta*. Biomolecules. 2021 Aug 15;11(8):1212.

52. Latorraca NR, Venkatakrishnan AJ, Dror RO. GPCR dynamics: structures in motion. Vol. 117, Chemical Reviews. American Chemical Society; 2017. p. 139–55.

53. Singh G, Inoue A, Gutkind JS, Russell RB, Raimondi F. PRECOG: PREdicting COupling probabilities of G-protein coupled receptors. Nucleic Acids Res. 2019 Jul 2;47(W1):W395–401.

54. Halton DW. Functional morphology of the platyhelminth nervous system. Vol. 113, Parasitology. Cambridge University Press; 1996.

55. Khan UW, Newmark PA. Somatic regulation of female germ cell regeneration and development in planarians. Cell Rep. 2022 Mar 15;38(11).

56. Terenina NB, Kreshchenko ND, Mochalova N V., Nefedova D, Voropaeva EL, Movsesyan SO, et al. The new data on the serotonin and FMRFamide localization in the nervous system of *Opisthorchis felineus* metacercaria. Acta Parasitol. 2020 Jun 1;65(2):361–74.

57. Roberts KM, Fitzpatrick PF. Mechanisms of tryptophan and tyrosine hydroxylase. IUBMB Life. 2013 Apr;65(4).

58. Patocka N, Ribeiro P. Characterization of a serotonin transporter in the parasitic flatworm, *Schistosoma mansoni*: cloning, expression and functional analysis. Mol Biochem Parasitol. 2007 Aug;154(2):125–33.

59. Richardson-Jones JW, Craige CP, Nguyen TH, Kung HF, Gardier AM, Dranovsky A, et al. Serotonin-1A autoreceptors are necessary and sufficient for the normal formation of circuits underlying innate anxiety. Journal of Neuroscience. 2011 Apr 20;31(16):6008– 18.

60. Mansour TE. The effect of serotonin and related compounds on the carbohydrate metabolism of the liver fluke, *Fasciola hepatica*. J Pharmacol Exp Ther. 1959 Jul;126(3):212–6.

61. Rahman MS, Mettrick DF, Podesta RB. *Schistosoma mansoni*: effects of *in vitro* serotonin (5-HT) on aerobic and anaerobic carbohydrate metabolism. Exp Parasitol. 1985 Aug;60(1):10–7.

62. Chen L, Huang S, Wu X, He W, Song M. Serotonin signalling in cancer: emerging mechanisms and therapeutic opportunities. Clin Transl Med. 2024 Jul 28;14(7).

63. Alenina N, Klempin F. The role of serotonin in adult hippocampal neurogenesis. Behavioural Brain Research. 2015 Jan;277:49–57.

64. Gaspar P, Cases O, Maroteaux L. The developmental role of serotonin: news from mouse molecular genetics. Nat Rev Neurosci. 2003 Dec;4(12):1002–12.

65. Tu Y, Yao S, Chen Q, Li W, Song Y, Zhang P. 5-Hydroxytryptamine activates a 5-HT/c-Myc/SLC6A4 signaling loop in non–small cell lung cancer. Biochim Biophys Acta Gen Subj. 2022 Apr 1;1866(4).

66. Dizeyi N, Hedlund P, Bjartell A, Tinzl M, Austild-Taskén K, Abrahamsson PA. Serotonin activates MAP kinase and PI3K/Akt signaling pathways in prostate cancer cell lines. Urologic Oncology: Seminars and Original Investigations. 2011 Jul;29(4):436–45.

67. Yu H, Qu T, Yang J, Dai Q. Serotonin acts through YAP to promote cell proliferation: mechanism and implication in colorectal cancer progression. Cell Communication and Signaling. 2023 Dec 1;21(1).

68. Du X, Wang T, Wang Z, Wu X, Gu Y, Huang Q, et al. 5-HT7 Receptor contributes to proliferation, migration and invasion in NSCLC cells. Onco Targets Ther. 2020 Mar;Volume 13:2139–51.

69. González A, Fazzino F, Castillo M, Mata S, Lima L. Serotonin, 5-HT1A serotonin receptors and proliferation of lymphocytes in major depression patients. Neuroimmunomodulation. 2007;14(1):8–15.

70. UniProt Consortium. UniProt: a worldwide hub of protein knowledge. Nucleic Acids Res. 2019 Jan 8;47(D1):D506–15.

71. Howe KL, Bolt BJ, Shafie M, Kersey P, Berriman M. WormBase ParaSite − a comprehensive resource for helminth genomics. Mol Biochem Parasitol. 2017 Jul;215:2–10.

72. Campos TDL, Young ND, Korhonen PK, Hall RS, Mangiola S, Lonie A, et al. Identification of G protein-coupled receptors in *Schistosoma haematobium* and *S. mansoni* by comparative genomics. Parasit Vectors. 2014 Dec 27;7(1):242.

73. Krogh A, Larsson B, von Heijne G, Sonnhammer ELL. Predicting transmembrane protein topology with a hidden markov model: application to complete genomes. Edited by F. Cohen. J Mol Biol. 2001 Jan;305(3):567–80.

74. Sayers EW, Beck J, Bolton EE, Bourexis D, Brister JR, Canese K, et al. Database resources of the National Center for Biotechnology Information. Nucleic Acids Res. 2021 Jan 8;49(D1):D10–7.

75. Jones P, Binns D, Chang HY, Fraser M, Li W, McAnulla C, et al. InterProScan 5: genome-scale protein function classification. Bioinformatics. 2014 May 1;30(9).

76. Sievers F, Wilm A, Dineen D, Gibson TJ, Karplus K, Li W, et al. Fast, scalable generation of high-quality protein multiple sequence alignments using Clustal Omega. Mol Syst Biol. 2011 Jan 11;7(1).

77. Tamura K, Stecher G, Kumar S. MEGA11: Molecular Evolutionary Genetics Analysis Version 11. Mol Biol Evol. 2021 Jun 25;38(7):3022–7.

78. Letunic I, Bork P. Interactive Tree Of Life (iTOL): an online tool for phylogenetic tree display and annotation. Bioinformatics. 2007 Jan 1;23(1):127–8.

79. Frickey T, Lupas A. CLANS: a Java application for visualizing protein families based on pairwise similarity. Bioinformatics. 2004 Dec 12;20(18):3702–4.

80. 80. Powell HR, Islam SA, David A, E Sternberg MJ. Phyre2.2: A community resource for template-based protein structure prediction [Internet]. Available from: https://esmatlas.com/

81. Schrödinger, LLC. The PyMOL molecular graphics system, version 2.5. New York: Schrödinger, LLC; 2021

82. Rosignoli S, Paiardini A. DockingPie: python-based tools for molecular docking workflows [Internet]. GitHub; [cited 2025 Jul 9]. Available from: https://github.com/forc-health-it/DockingPie

83. Burley S, Berman H, Kleywegt J, Nakamura H, Velankar S. Protein Data Bank (PDB): the single global macromolecular structure archive. Protein crystallography: methods and protocols. 2017;627–41.

84. McVeigh P, McCammick EM, McCusker P, Morphew RM, Mousley A, Abidi A,et al. RNAi Dynamics in juvenile Fasciola spp. liver flukes reveals the persistence of gene silencing in vitro. PLoS Negl Trop Dis. 2014 Sep 25;8(9):e3185.

85. McCusker P, McVeigh P, Rathinasamy V, Toet H, McCammick E, O’Connor A, et al. Stimulating neoblast-like cell proliferation in juvenile Fasciola hepatica supports growth and progression towards the adult phenotype in vitro. PLoS Negl Trop Dis. 2016 Sep 13;10(9).

86. Armstrong R, Marks NJ, Geary TG, Harrington J, Selzer PM, Maule AG. Wnt/β-catenin signalling underpins juvenile Fasciola hepatica growth and development. PLoS Pathog 2025 7;21(2):e1012562.

87. Jing L. Preparation of torula yeast RNA for hybe solutions [Internet]. Available from: http://www.bio-protocol.org/e180

88. King RS, Newmark PA. In situ hybridization protocol for enhanced detection of gene expression in the planarian *Schmidtea mediterranea*. BMC Dev Biol. 2013;13(1).

89. Schneider CA, Rasband WS, Eliceiri KW. NIH Image to ImageJ: 25 years of image analysis. Nat Methods. 2012 Jul 28;9(7).

90. Nussbaum-Krammer CI, Neto MF, Brielmann RM, Pedersen JS, Morimoto RI. Investigating the spreading and toxicity of prion-like proteins using the metazoan model organism *C. elegans*. Journal of Visualized Experiments. 2015 Jan 8;(95).

91. McCusker P, Hussain W, McVeigh P, McCammick E, Clarke NG, Robb E, et al. RNA interference dynamics in juvenile *Fasciola hepatica* are altered during in vitro growth and development. Int J Parasitol Drugs Drug Resist. 2020 Dec;14.

92. Pfaffl MW. A new mathematical model for relative quantification in real-time RT-PCR. Nucleic Acids Res. 2001 May 1;29(9).

